# Population growth affects intrinsic and extrinsic noise in gene expression

**DOI:** 10.1101/362368

**Authors:** Philipp Thomas

## Abstract

Clonal cells of exponentially growing populations vary substantially from cell to cell. The main drivers of this heterogeneity are the population dynamics and stochasticity in the intracellular reactions, which are commonly studied separately. Here we develop an agent-based framework that allows tracking of the biochemical dynamics in every single cell of a growing population that accounts for both of these factors. Apart from the common intrinsic variability of the biochemical reactions, the framework also predicts extrinsic noise arising from fluctuations in the histories of cells without the need to introduce fluctuating rate constants. Instead, these extrinsic fluctuations are explained by cell cycle fluctuations and differences in cell age, which are ubiquitously observed in growing populations. We give explicit formulas to quantify mean molecule numbers, intrinsic and extrinsic noise statistics as measured in two-colour experiments. We find that these statistics may differ significantly depending on the experimental setup used to observe the cells. We illustrate this fact using (i) averages over an isolated cell lineage tracked over many generations as observed in the mother machine, (ii) snapshots of a growing population with known cell ages as recorded in time-lapse microscopy, and (iii) snapshots of unknown cell ages as measured from static images. Our integrated approach applies to arbitrary biochemical networks and generation time distributions. By employing models of stochastic gene expression and feedback regulation, we elucidate that isolated lineages, as compared to snapshot data, can significantly overestimate the mean number of molecules, overestimate extrinsic noise but underestimate intrinsic noise and have qualitatively different sensitivities to cell cycle fluctuations.

## I Introduction

The behaviour of clonal cells varies substantially from cell to cell and over time^1,2^. Identifying the sources of these fluctuations can help us to understand how clonal cells diversify their responses and to reveal the function of gene circuits and signalling networks. For their quantification, it is often convenient to break down the experimentally observed variability into functional components. Commonly one wishes to separate fluctuations inherent in the circuit dynamics itself, called *intrinsic noise*, from fluctuations that arise from embedding the circuit in the environment of the cell, called *extrinsic noise*.

A possible resolution to this problem is to place and simultaneously measure a second independent copy of the circuit in the cell, as has been done in *E. coli*^1^, yeast^3^, mammalian cells^4^ and plants^5^. The difference between the two circuit copies measures the intrinsic noise whereas their correlations measure the extrinsic noise component. Intrinsic noise arises from the random nature of the involved biochemical reactions. Extrinsic noise originates from factors affecting both circuits in the same way. These can, for instance, be modelled by reaction rates that fluctuate between cells or over time due to shared resources, promoter architecture or upstream pathways. Such sources of extrinsic noise have been studied extensively in the literature^6–14^.

A less commonly studied but equally important source of extrinsic noise is the population dynamics^15^. Since intracellular molecule numbers must double over the cell division cycle, a two-fold variation of expression levels is expected from cell proliferation alone. Moreover, the cell cycle itself is subject to tremendous variation providing an additional source of extrinsic variability. For example, generation times in *Escherichia coli*^16^, budding yeast^17^ and mammalian cells^18^ vary about 30 – 50%. These sources should therefore prevail in growing cells, populations and tissues.

Modelling approaches for understanding the effects of the cell cycle on gene expression noise are only recently being developed^19–24^. These studies are often restricted to a single isolated cell observed over successive cell divisions and measuring variability over time, similar to a lineage in the mother machine^25^. Many experiments, however, report cell-to-cell variability across snapshots of an exponentially growing cell population. These approaches either use time-lapse microscopy^26,27^ or analyse snapshots with distributed cell ages as observed in flow cytometry, smFISH or similar techniques^28–30^.

Recent studies elucidated that population snapshots and lineages can significantly deviate from each other in their statistics^24,31^. To-date, however, there exists no general analytical framework with which to quantify the gene expression fluctuations in populations. We are thus lacking the means with which to understand, compare and integrate the knowledge from different experiments such as mother machines, time-lapse microscopy or fixed-cell images. Agent-based approaches allowing to track the expression levels of each individual cell in a growing population are ideally suited to address this issue.

In this manuscript, we develop such an approach to characterise the statistics of biochemical reaction networks in a growing and dividing cell population. In this framework, an agent is represented by a cell whose biochemical decomposition changes due to stochastic reaction kinetics and cell divisions. In Sec. II A we show how to analytically characterise the joint distribution of cell age and molecule content per cell in a snapshot of the population. We then, in Sec. II B, derive the exact moment equations of this model.

Since stochastic models are rarely solvable, we propose an analytically tractable approximation to mean and covariances in Sec. III A. Intrinsic and extrinsic noise sources as they are measured using two-reporter systems are in-built in the agent-based approach, and we explain how to decompose the apparent noise into the respective components. We further elaborate on the decomposition in cases where the cell age is unknown, a situation commonly encountered when analysing data from population snapshots or flow cytometry. We demonstrate how to practically compute the noise decomposition in Sec. III B, illustrate the results using a simple two-reporter system, and study how circuit dynamics can be tuned to suppress either intrinsic or extrinsic fluctuations.

## II Methods

We model the dynamics of a dividing population of cell agents. The state of each cell is given by its age and the number of intracellular molecules, which evolve from birth to division. After cell division, the mother’s molecules are inherited by the two daughters through stochastic partitioning of molecules. Cell divisions occur asynchronously in the population because cells divide at random times. In consequence, cell ages and molecule numbers are heterogeneous in the population. Fig. 1a illustrates the resulting branching process whose final state is a snapshot of the cell population.

**FIG. 1.**
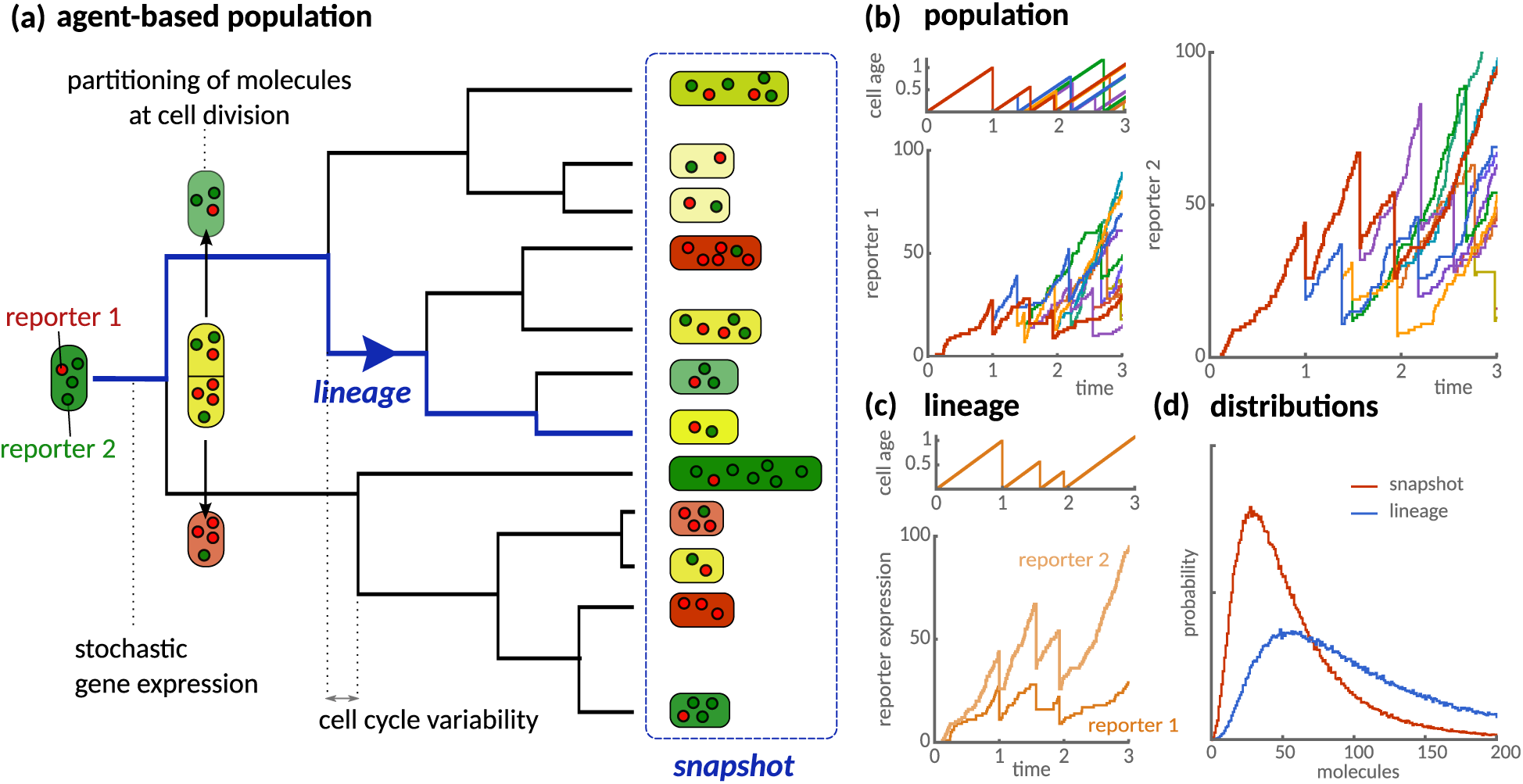
Agent-based model of clonal population dynamics with stochastic gene expression and cell cycle variability. (**a**) Illustration of a growing population as a stochastic branching process with stochastic interdivision times. Each cell expresses two identical but noninteracting reporters (green and red) that are partitioned randomly at cell division. Red and green cells express more molecules of either reporter, which indicates intrinsic variability between cells. Yellow cells express similar levels of reporter molecules, but vary in their absolute amounts at different cell cycle stages, which constitutes extrinsic variability. A snapshot of the population (blue dashed box) quantifies the cell-to-cell variability across the population. A lineage (red path) quantifies variability over time and tracks an isolated cell over successive cell divisions by randomly selecting one of the daughter cells. (**b**) Simulated trajectories of cell age and stochastic protein expression of two identical reporters on a branched tree. Line colour indicates reporter expression in the same cell. (**c**) Cell age and reporter expression of an isolated cell lineage. (**d**) Comparison of distributions obtained from lineages and population snapshots. Simulations of the reactions (21) assume *k*_0_ = 10, *k_m_* = 1, *k_s_* = 10 for each reporter and lognormal-distributed division times with unit mean and standard deviation.

The seminal experiment by *Elowitz et al*. identified the sources of cell-to-cell variation using snapshots of cells expressing green and red fluorescent reporters^1^. Reporters expressed at different levels appear either red or green, a signature of intrinsic noise. Cells with similar reporter levels appear yellow but with variable intensities, a signature of extrinsic noise. In our model (Fig. 1a), similar effects are observed since stochasticity in biochemical reactions and partitioning of molecules at division account for intrinsic variation across the population. Cell age and variability in division timing provide a source of extrinsic noise (Fig. 1b). In contrast to the population-view, the dynamics of isolated cells can also be tracked over time, which we will refer to as lineage (Fig. 1c), which corresponds to a random path in the tree. These statistics can differ significantly from population snapshots (Fig. 1d). To develop a quantitative understanding of these effects, we begin with deriving analytical framework to quantify these populations.

### A Agent-based framework for stochastic biochemical kinetics in growing cell populations

To each cell we associate an age *τ* that counts the time since its last division and a number of set of biochemical species *X*_1_, *X*_2_, *…*, *X_NS_* present in amounts *x* = (*x*_1_, *x*_2_, …, *x_Ns_*). These species interact via a network of *R* intracellular biochemical reactions of the type

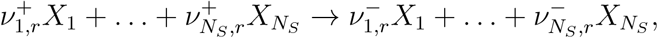

where 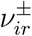 are the stoichiometric coefficients and *r* = 1, …, *R*. In the following, we outline the master equation that allows to analytically study these networks in an agent-based context.

#### 1 Master equation for the agent-based population

The state of the overall cell population can be characterised by the snapshot density *n*(*τ, x, t*) that counts the number of cells at time *t* with age between *τ* and *τ* + d*τ* and molecule counts *x*. Accordingly, the total number of cells in the population is given by

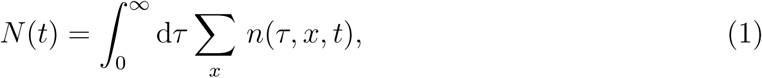

where the summation is over all possible molecule number configurations *x*.

We assume that cells divide with an age-dependent rate *γ*(*τ*), which is related to the interdivision time distribution *φ* (*τ_d_*) via

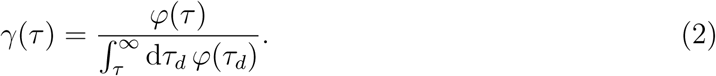

The snapshot density then evolves due to age-progression of cells, cell divisions and the change in their molecular decomposition due to biochemical reactions

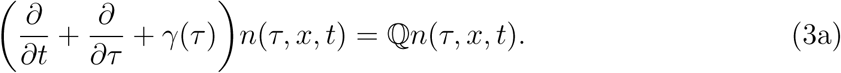

Here, the change in the molecule numbers per cell is expressed by the transition matrix ℚ acting as 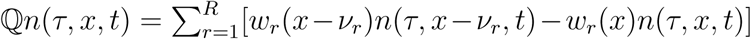, where 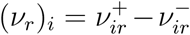 is the stochiometric vector of the *r^th^* reaction. Cell birth is described by the boundary condition

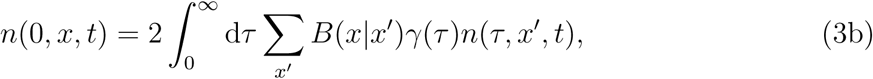

whereby the mother cell is replaced with two daughter cells of zero age with its molecules being partitioned between them. The division kernel *B*(*x|x′*) is the probability of partitioning the molecule numbers *x′* to *x* of any daughter cell and is given by

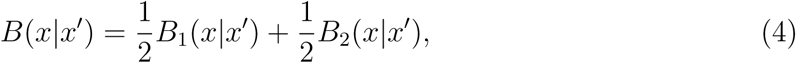

where *B*_1_ and *B*_2_ are the marginal probabilities for the two daughter cells to inherit *x* of the mother cell’s molecules *x′*. Importantly, if the total amount is conserved in the division, we have *B*_2_(*x|x′*) = *B*_1_(*x′*− *x|x′*). Thus each cell inherits an equal amount of molecules *E_B_*[*x|x′*] = *x′/*2, because we do not distinguish the daughters.

Since resolving the time-evolution of the snapshot density is a formidable task, we concentrate on the long-term evolution of Eq. (3), which describes the exponential growth phase or balanced growth condition. In this limit, the total number of cells grows exponentially *N*(*t*) ~ *N*_0_*e^λt^* with rate *λ* and the fraction of cells with a certain cell age and molecule content is constant

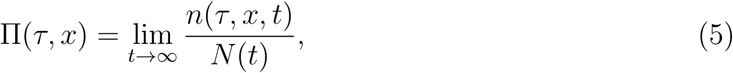

due to the balance between cell births, divisions and the biochemical reactions. In the following, we summarise how to characterise this distribution analytically.

#### 2 Age-distribution and population growth rate

The fraction of cells with the same age in a snapshot is given by the age-distribution, which follows^31^

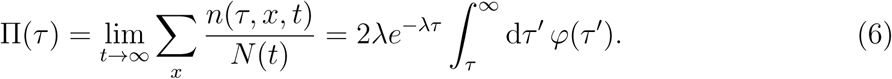

The distribution *φ* characterises the interdivision times

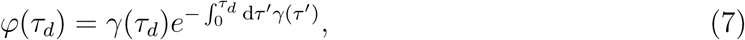

as also seen from Eq. (2). The age distribution, Eq. (6), depends on the population growth rate *λ* that is uniquely determined by the Euler-Lotka equation

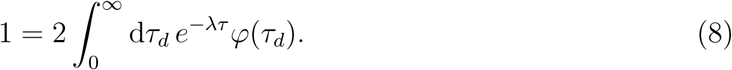

The above equations constitute the fundamental age-structure of microbial populations, which has been verified in experiments^16,32,33^.

#### 3 Distribution of molecules for cells of the same age

We consider the total number of cells with age *τ* and molecule count *x* divided by the number of cells at that age. This conditional probability quantifies the likelihood of observing *x* molecules in a cell of age *τ* and is given by

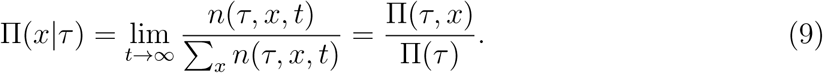

It can be verified^31^ that Π(*x|τ*) obeys

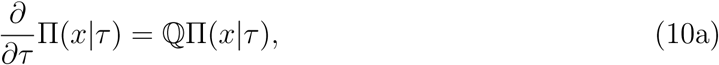

which is similar to the chemical master equation (with time replaced by cell age). An important difference though, is that it has to be solved subject to the boundary condition

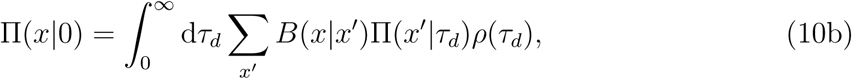

which accounts for the cell divisions. The distribution under the integral

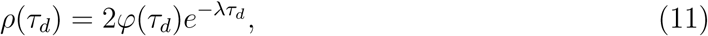

is the interdivision time distribution in the population^32,34^. The distribution describes the interdivision times of cells with completed cell divisions and depends explicitly on the population growth rate *λ*.

#### 4 Comparison with the lineage framework

A lineage tracks a single of the population over successive cell divisions. In the long term its evolution also approaches a stable distribution, which we denote by *π*(*τ, x*). The molecule number distribution for cells of the same age in a lineage is given by *π*(*x|τ*) = *π*(*x, τ*)/*π*(*τ*) and satisfies

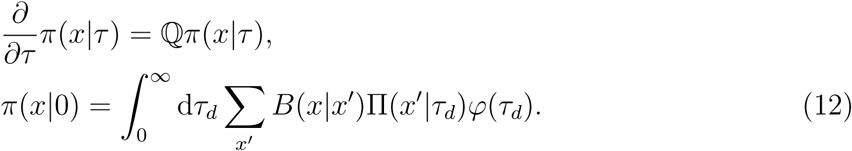

By comparing the above equations with Eqs. (10), we notice that this distribution is obtained by substituting the division time distribution *ρ* by *φ*. Thus cells of the same age can be analysed using a unified framework whether in populations or lineages. The age-distribution in a lineage, however, which is

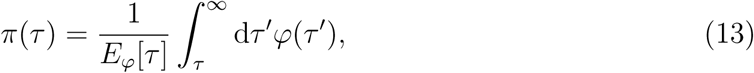

differs significantly from the population, Eq. (6).

### B Moment equations for the agent-based model

In many practical situations, solving for the full distribution is infeasible. Summary statistics such as means and variances, which we will focus on in the following, present convenient alternatives as they are more amenable to analysis.

#### 1 Exact moment equations for cells of the same age

In brief, the moment equations are obtained by multiplying Eq. (10a) by *x* or *xx^T^* and summing over all possible states. The results for the first and second moments are

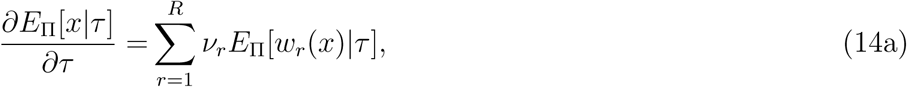

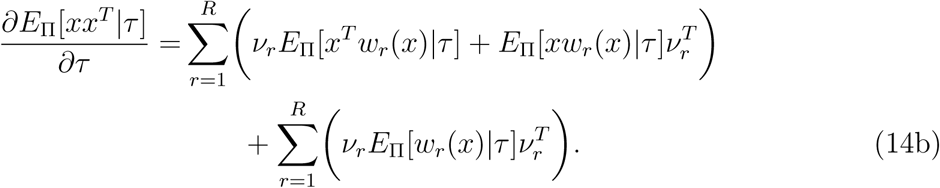

Interestingly, these are the same moment equations that appear in the study of systems without age-dependence (with age being replaced by the observational time). The key difference is the boundary condition subject to which the moment equations have to be solved. These conditions follow from Eq. (10b) and the conservation of molecules in Eq. (4), which implies *E_B_*[*x|x′*] = *x′/*2. They read

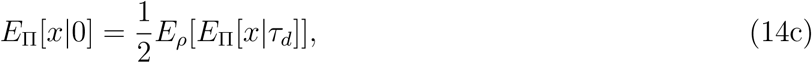

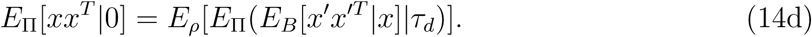

The first condition states that, on average, molecule numbers need to double over one cell cycle. The second condition relates the second moments to the partitioning of molecules described by the division kernel, Eq. 4.

#### 2 Exact moment equations for cells of unknown age

We now consider the snapshot moments of molecule numbers irrespective of age. *E*_Π_[*x*] = *E*_Π_[*E*_Π_[*x|τ*]] and *E*_Π_[*xx^T^*] = *E*_Π_[*E*_Π_[*xx^T^* |*τ*]] are obtained by multiplying Eq. (14) with Π (*τ*) and performing the integration. For this purpose, consider the expected value of a function *f*(*τ*) with respect to the age distribution, which satisfies

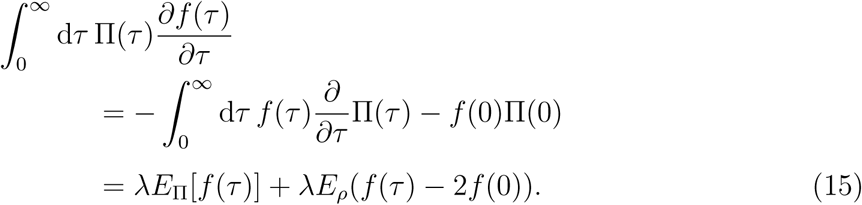

In the first line, we integrate by parts assuming lim_*τ*→∞_ *f*(*τ*) Π(*τ*) = 0, and in the second line we substituted Eq. (6) for Π(*τ*) and performed the derivative. The first term captures the effect of dilution, while the second term describes discrete changes during cell division.

Setting now *f*(*τ*) = *E*_Π_[*x|τ*] in (15) and combing the result with Eq. (14a) and the boundary conditions (14c), we find an equation for the mean number of molecules in the population,

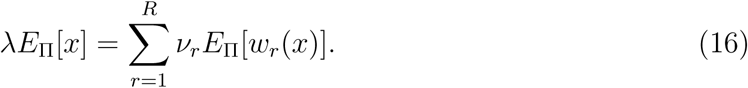

Similarly, using Eq. (15) with *f*(*τ*) = *E*_Π_[*xx^T^ |τ*], Eq. (14b) and (14d), the equation for the second moment becomes

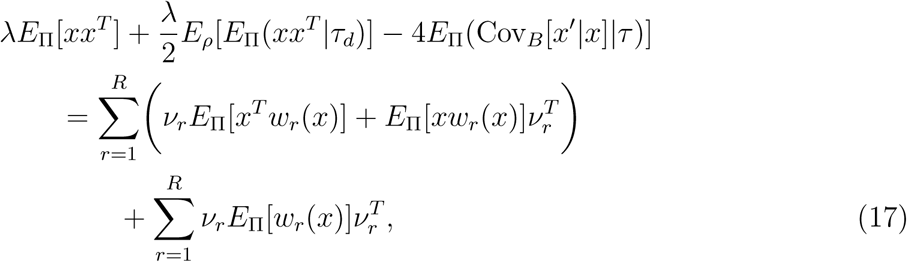

where the left hand side depends explicitly on the division-time distribution *ρ*. Obviously, these equations cannot be solved in general, not only because the hierarchy of moments is not closed but also because they depend on moments for cells of known age. The conditions for which these equations are closed and can be solved exactly are discussed in SI V C.

## III Results

### A Decomposing noise into intrinsic and extrinsic contributions

To circumvent the moment-closure problem, we employ the linear noise approximation to decompose the noise into intrinsic and extrinsic components (see^35,36^ for details of the approximation). In brief, the approximation assumes Gaussian fluctuations and provides closed-form expressions for the mean molecule numbers and their covariances. Writing short Cov_Π_[*x|τ*] = *E*_Π_ [*xx^T^ |τ*] – *E*_Π_ [*x|τ*]*E*_Π_ [*x^T^ |τ*] for the covariance matrix, the result is

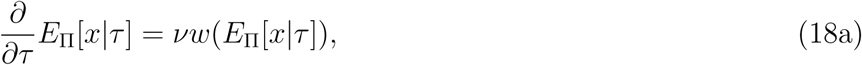

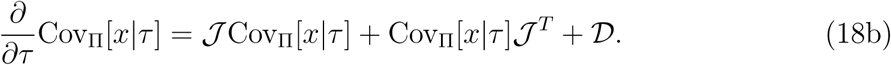

where the Jacobian 𝒯 and the diffusion matrix 𝒟 are defined as

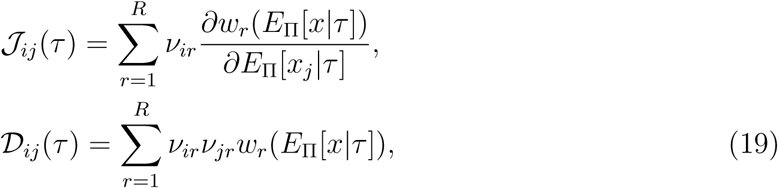

which depend on cell age through the mean molecule numbers *E*_Π_[*x|τ*]. Comparison of Eqs. (18) with (14) shows that these equations are exact whenever the propensities are linear in the molecule numbers. In all other cases, we consider these as an approximation valid in the limit of large molecule numbers.

Next, we cast the boundary condition (14d) in terms of the covariance matrix Cov_Π_[*x|τ*], which leads to

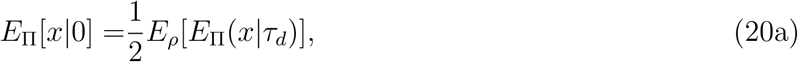

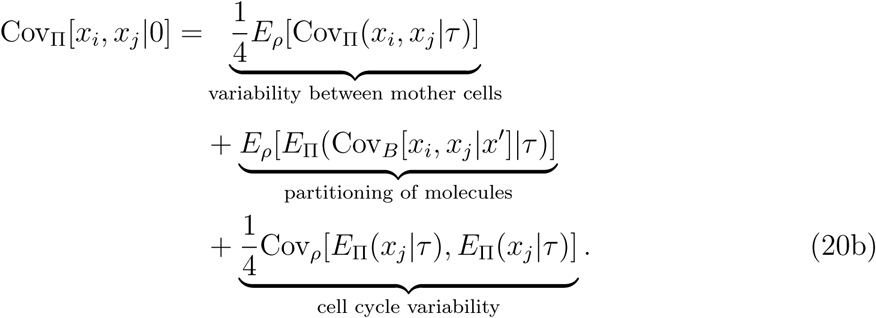

The first term is the contribution due to fluctuations in the number of molecules before division. The second term denotes the variation due to random partitioning of molecules at cell division, while the third contribution stems from differences in the molecule numbers due to different cell cycle lengths. We note that Eqs. (20b) themselves do not constitute a noise decomposition since these contribute do not propagate independently. Instead, they represent the sources of cell-to-cell variability for the two daughter cells.

#### 1 Noise decomposition using dual reporter systems

To investigate how the different sources of variations affect biochemical reaction dynamics, we consider the synthesis and degradation of mRNA molecules and translation into proteins

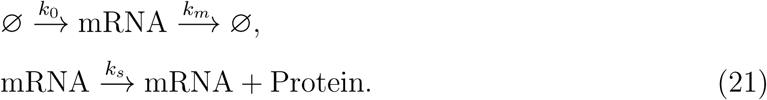

We do not account for protein degradation explicitly in this model since stable proteins are diluted in the population by cell division, the effect of which we will study in the following. For simplicity, we assume that mRNA degradation is faster than the population growth such that the reactions can be approximated by a single reaction synthesising proteins in stochastic bursts. At the same time, for the purpose of the noise decomposition, we consider an additional, identical copy of the same circuit in the cell

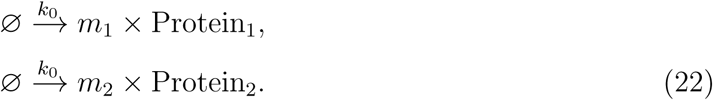

The stochastic burst size of the first and second copy are denoted by *m*_1_ and *m*_2_, respectively, and both follow a geometric distribution with mean *b* = *k_s_/k_m_* (see Ref.^37^ and SI V D for details of the burst approximation).

##### a. Mean number of proteins

Since the two reporter proteins are expressed identically in the cell, their mean expression levels must be the same. Denoting the protein numbers of the two reporters by *x*_1_ and *x*_2_, we have *E*_Π_[*x*_1_*|τ*] = *E*_Π_[*x*_2_*|τ*]. The rate equation (18a) for the average number of proteins for a cell of given age becomes

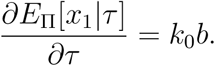

The solution that respects the boundary condition (20a) is

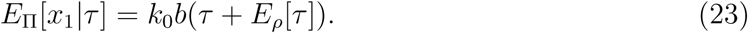

The number of proteins inherited after cell division (*τ* = 0) is thus *k*_0_*bE_ρ_*[*τ*], which depends on the mean division time *E_ρ_*[*τ*] in the population.

##### b. Separating noise into intrinsic and extrinsic components

For identical two-reporter systems, the overall variance can be decomposed into intrinsic and extrinsic components as follows

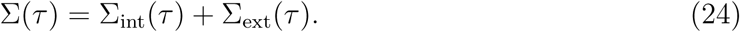

The individual contributions can be quantified using^1^

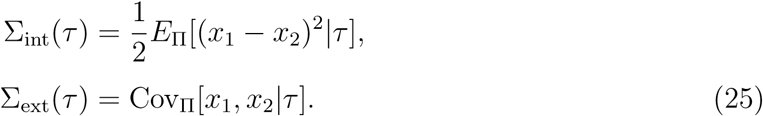

Since these components are measured in the same cell, they also account for the correct history dependence^7,9,38^.

The variance of intrinsic and extrinsic fluctuations follows from using Eqs. (25) in (18b) and rearranging, which leads to

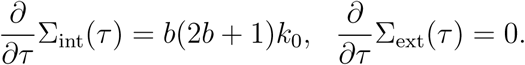

Its solution is obtained by straight-forward integration and is given by Σ_int_(*τ*) = Σ_int_(0) + *b*(2*b* + 1)*k*_0_*τ* and Σ_ext_(*τ*) = Σ_ext_(0). To fix the boundary condition (20b), we assume that each molecule of the mother cell being partitioned with equal probability between the two daughter cells. In this case, the division kernel in Eq. (4) is binomial with covariance Cov*_B_*[*x_i_, x_j_|x′*] = δ*_ij_x′_i_/*4. We then find that 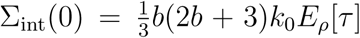 and Σ_ext_(0) = 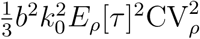, and finally

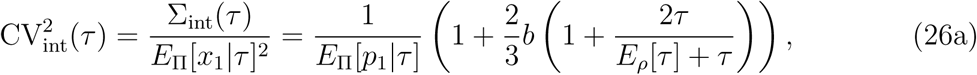

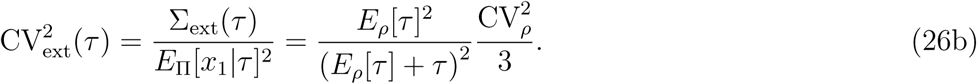

The coefficients of variations quantify the size of fluctuations relative to the mean. The result confirms the intuition that intrinsic noise decreases with the mean number of molecules. The extrinsic noise component, however, reflects the variations in cell cycle duration 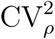 that are transmitted to the protein levels.

##### c. Snapshots display higher intrinsic but lower extrinsic noise levels than lineages

Next, we compare the statistics of snapshots of a growing population with the one of a lineage of an isolated cell over time. As explained in Sec. II A 4, we obtain the lineage statistic by substituting the division time distribution *ρ* for *φ* in Eqs. (23), (26a) and (26b).

Interestingly, the deviations between these two statistics is apparent even on the mean level. To see this, we notice that the mean number of molecules, Eq. (23), increases with the duration of the cell cycle. It is well known that the cell cycle time is longer when averaged over single cells than for cells in the population^16^ *E_ρ_*[*τ*] ≤ *E_φ_*[*τ*]. An intuitive explanation of this fact is that fast dividing cells are over-represented in the population. It hence follows from Eq. (23) that the expected number of molecules is lower in populations compared to lineages, no matter what the division time distribution is.

In Fig. 2a, we compare the total noise 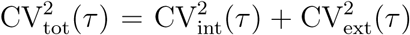 for gamma and log-normal distributed interdivision times. In both cases, we observe that the noise exhibits a maximum for low cell cycle variability. With increasing cell cycle variability, we find that the maximum flattens in the lineage but not in snapshot statistics. Albeit the two statistics are collected from different samples of the same population, snapshots are more noisy than lineages in both cases. To understand this noise propagation, we decompose the total noise into intrinsic and extrinsic components via Eqs. (26a) and (26b). We observe that intrinsic noise in snapshots increases with cell cycle variability (Fig. 2b) while it is significantly lower in lineages and independent of these fluctuations, which is consistent with lower expression levels in snapshots.

**FIG. 2.**
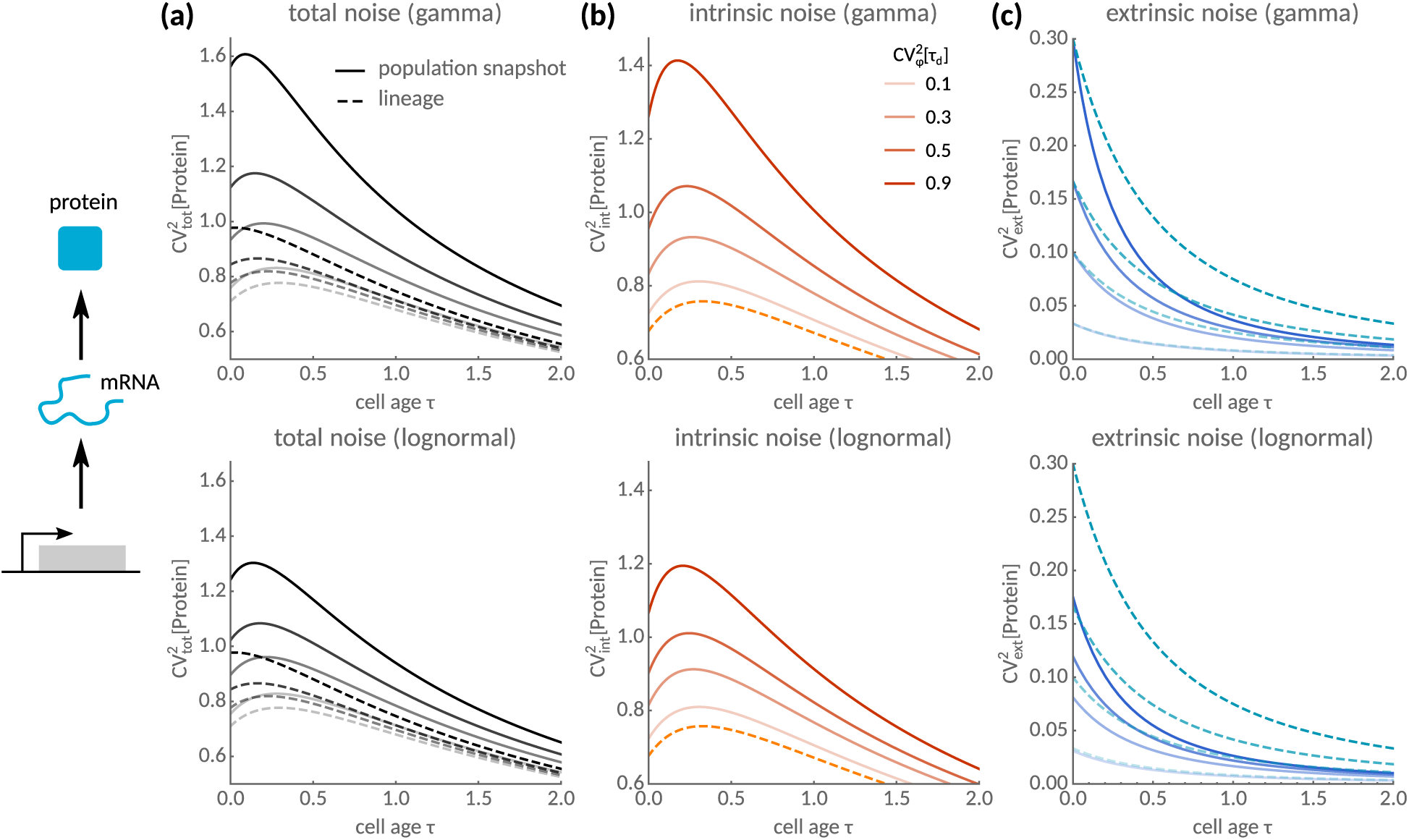
Intrinsic and extrinsic noise propagation over the cell cycle. (**a**) Total noise as a function of cell age *τ* with gamma (top) and log-normal-distributed (bottom) interdivision times. Population snapshot statistics (solid) are compared to lineages (dashed lines). Noise is monotonic for large cell cycle fluctuations CV^2^[*τ_d_*] in lineages but not in snapshots. (**b**) Intrinsic noise peaks as a function of cell age and increases with cell cycle fluctuations in populations but not in lineages. (**c**) Extrinsic noise is lower in the population than in lineages. Parameters are *k*_0_ = 1, *b* = 100 and division time distributions assume unit mean.

Fig. 2a also reveals a non-monotonic dependence of the intrinsic noise component on cell age. To explain this phenomenon, we notice that intrinsic noise, Eq. (26a), increases with cell age due to an increase in the Fano factor. For older cells, however, intrinsic noise decreases with age as these cells express higher protein levels. Combining these findings explains the noise maximum at a well-defined cell age. By maximising Eq. (26a) over all possible cell ages, the age at which noise peaks is 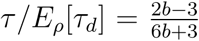 whenever *b* > 3/2. This ratio only depends on the burst size *b* and approaches 1/3 of the mean cell cycle time for large *b*. By contrast, we find that extrinsic noise is lower and decays slower over the cell cycle in snapshots than in lineages (Fig. 2c). We conclude that lineage statistics may significantly underestimate intrinsic heterogeneity but overestimate extrinsic noise in the population. In the next subsection, we extend this method to general stochastic reaction networks.

#### 2 General decomposition for cells of the same age

We now generalise the decomposition to two-reporter systems involving an arbitrary network of biochemical reactions. As before, we assume that the two copies of our network with molecule numbers *x*_1_ and *x*_2_ do not interact and involve the same type reactions, and therefore they have the same mean expression level. The covariance of the two-reporter system is

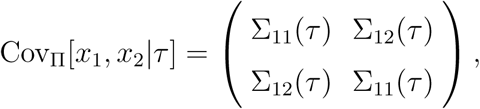

whose individual components satisfy

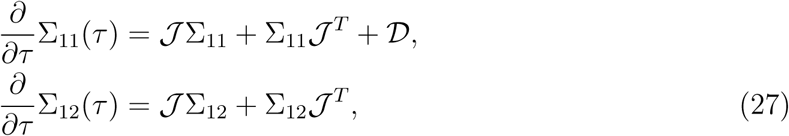

according to Eqs. (18b).

The intrinsic and extrinsic noise components can be expressed via

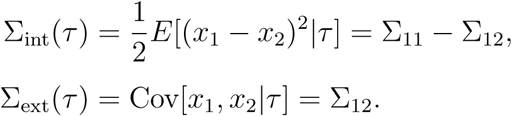

Since the covariances obey the linear equations (27), the two noise contributions evolve independently. In particular, the intrinsic and extrinsic covariances satisfy

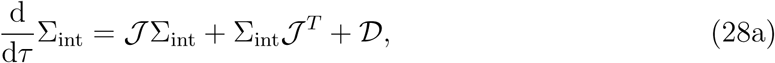

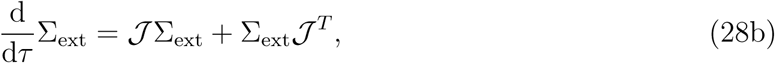

where only the intrinsic component involves the biochemical noise from the intracellular reactions through the diffusion matrix 𝒟.

Assuming again binomial partitioning with covariance Cov*_B_*[*x_i_, x_j_|x′*] = δ*_ij_x′_i_/*4, allows us to split the boundary condition (20b) according to Eq. (24). The results are two independent conditions

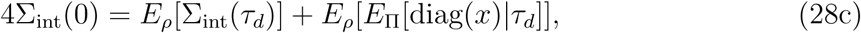

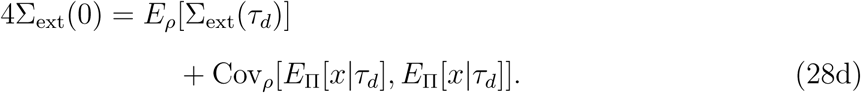

The noise decomposition is fully specified by the mean number of molecules for cells of the same age, the Jacobian 𝒯 of the corresponding rate equations, the diffusion matrix 𝒟 (see Eqs. (19)) and the distribution of interdivision times in the population *ρ* (see Eq. (11)). Importantly, Eq. (28c) shows that partitioning is a noise source to intrinsic fluctuations, while Eq. (28d) shows that cell cycle variations contribute to extrinsic fluctuations. We conclude that conditioning on the cell cycle position is not enough to eliminate all extrinsic noise. Before we continue, we note that other types of partitioning, such as asymmetric cell division, can be easily incorporated into the framework but using a different form of Cov*_B_*[*x_i_, x_j_|x′*]^39^.

#### 3 General decomposition for cells of unknown age

An obstacle for applying this decomposition in practice is that in many situation the cell age is not known, and this is especially true for population snapshots. For this reason, the mean of the molecule number has to be averaged over all possible cell ages

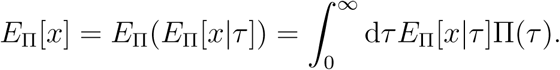

Similarly, we use the law of total variance to decompose the snapshot-variance as

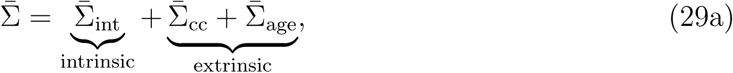

with

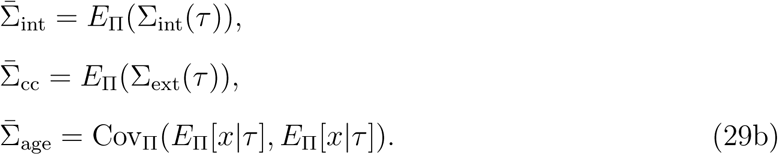

The first term in Eq. (29a) is the intrinsic variance measured across a population, the second term is the extrinsic variance transmitted from cell cycle variations, and the third term is another extrinsic component that comes from averaging over cells of different ages. The total extrinsic noise, which is measured in a two-reporter system, is the sum of the second and third term. The practical use of this noise decomposition is demonstrated in the following section.

##### B Practical computation of the noise decomposition and applications

Finally, we apply the noise decomposition to analyse snapshots in which the age of individual cells is not known. While the decomposition can be carried out exactly for linear reaction networks, we also outline a numerical method with which the decomposition can be carried out efficiently for complex nonlinear networks as we demonstrate for a protein that regulates its own expression.

#### 1 Decomposition for linear reactions in cells of unknown age

For linear reaction networks in which the propensities are linear functions of the molecule numbers. This dependence allows to average the statistics exactly over all cell ages. Thus from Eq. (16), we obtain the rate equations

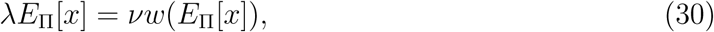

since *E*_Π_[*w*(*x*)] = *w*(*E*_Π_[*x*]). These equations coincide precisely with the steady state of the traditional deterministic rate equations including an effective dilution term proportional to the population growth rate *λ*.

Averaging Eq. (28a) over all ages and accounting for the boundary terms using Eq. (15), the intrinsic variance becomes

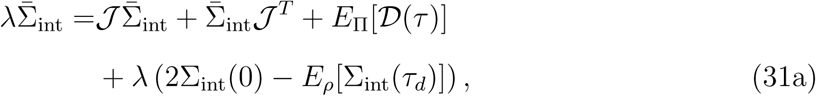

where the Jacobian 𝒯 is assumed to be independent of cell age. Similarly, averaging Eq. (28b) the extrinsic variance transmitted from cell cycle fluctuations is obtained as

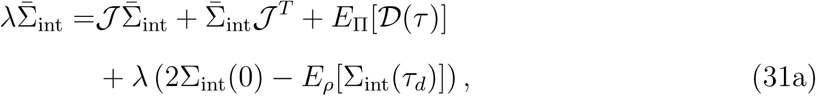

Similarly, an equation for 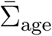 can be derived (see SI V B for details), which reads

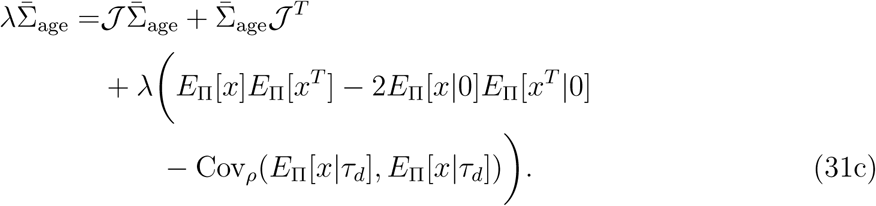

This decomposition exactly characterises the variability of linear intracellular reaction networks across snapshots.

##### a. Application to stochastic reporter expression

We return to the two-reporter system (22) and apply the decomposition developed in the previous section. From Eq. (30), we find that the mean molecule number is given by

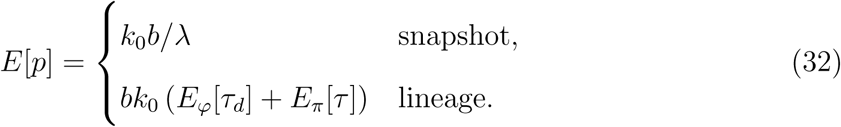

Note that the lineage mean follows from integrating Eq. (23) with *φ* instead of *ρ* against the lineage age-distribution (13). Interestingly, only the population mean agrees with the intuition in which the ratio of synthesis and dilution rates yields the steady state levels. However, both averages depend implicitly on the cell cycle variability through the average age *E_π_*[*τ*] or the population growth rate *λ*, respectively. In Fig. 3a, we show that molecule numbers in the lineage increase with cell cycle variability while they decrease in the snapshot statistic under the same conditions. These quantities thus exhibit opposite sensitivities to cell cycle variability.

**FIG. 3.**
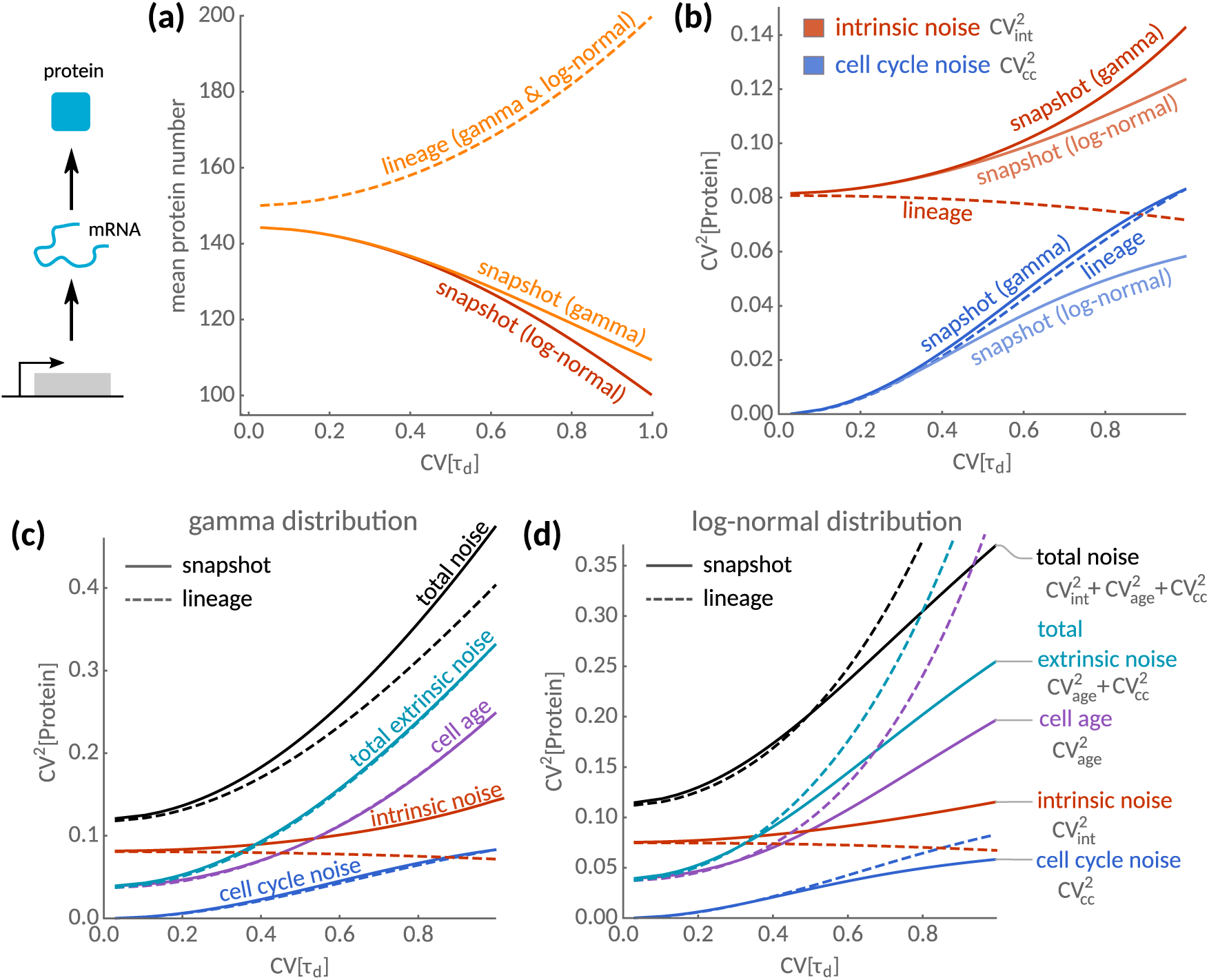
Statistics of population snapshots and isolated lineages for cells of unknown age. (**a**) Mean protein number as a function of the cell cycle variations CV[*τ_d_*] in lineages (dashed) and snapshots (solid lines). For lineages, the mean protein number increases with cell cycle variability and is independent of the division time distribution. In snapshots, the mean decreases with cell cycle variability with a rate that depends on higher moments of the distribution. The predictions for gamma- and log-normal distributed interdivision times is shown. (**b**) Sensitivity of intrinsic and extrinsic noise sources to cell cycle fluctuations. Intrinsic noise (red lines) increases in lineages but decreases in snapshots consistent with the dependence of the respective means shown in (a). The transmitted cell cycle noise (blue lines) shows a similar dependence on cell-cycle variability in lineages and snapshots for the gamma-distribution, but is lower in snapshots for the log-normal distribution. (**c**) Total noise (black lines) broken down into individual noise components for the gamma-distribution. Transmitted cell cycle noise and the uncertainty due to distributed cell ages (purple lines) contribute to the total extrinsic noise (teal). (**d**) The corresponding break-down for the log-normal distribution. Parameters are *k*_0_ = 10 and *b* = 10 and *E_φ_*[*τ_d_*] = 1.

Next, we explore the noise properties of the reporter system using the decomposition (31). We find that the contributions of intrinsic noise are

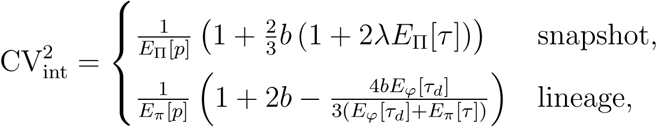

which is inversely proportional to the mean number of proteins. The contribution of extrinsic noise due to stochasticity in cell cycle duration is

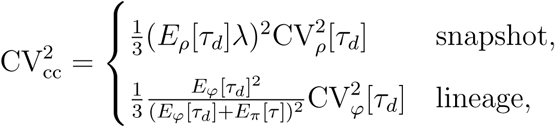

and the one due to the unknown cell age is

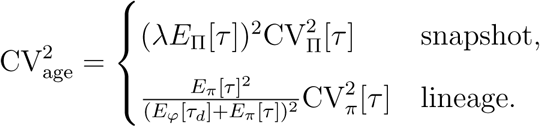

The noise decomposition crucially depends on the population growth rate *λ*, while in lineages it depends on the corresponding average cell age *E_π_*[*τ*]. More specifically, 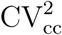 and 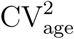 depend the variations in the age and interdivision time distributions, which are generally different in lineages and population. We illustrate this dependence using the analytical decompositions for two different interdivision-time distributions with the same mean and variance.

For both the gamma and the log-normal distribution, intrinsic noise (red lines, Fig. 3b) exhibits opposite sensitivities on cell cycle variability comparing lineage (dashed) and snapshot statistics (solid). This observation is explained by smaller mean expression levels in snapshots (cf. Fig. 3a) because intrinsic noise is expected to scale inversely with the mean molecule number. For the gamma distribution, the extrinsic noise transmitted from cell cycle variations (blue lines, Fig. 3b) is (almost) identical for these measures. For the log-normal distribution, however, extrinsic noise in the lineage is smaller than in the snapshot. Interestingly, we find that the total noise is higher in snapshots than in lineages for the gamma distribution (black lines, Fig. 3c), while this not true for the log-normal distribution and large cell cycle variability (black lines, Fig. 3d).

In developed network models, the extrinsic components will also depend on the biochemical properties of the network. We demonstrate this analytically in SI V D when the protein is also subject to degradation, which reveals intricate noise patterns. A straight-forward approach for the noise decomposition in complex biochemical network is given in the following section.

#### 2 Decomposition for nonlinear reaction networks

For nonlinear reaction networks, it is generally difficult to carry out the noise decomposition analytically. This is because the statistics of known and unkown cell age are intricately coupled and can be solved simultaneously only in simple cases. An efficient and generally applicable procedure to compute the numerical noise decomposition is the following:

1. Calculate the population growth rate using Eq. (8).
2. Solve for the statistics of cells of the same age, Eq. (28a) and (28b) and use the shooting method to match the boundary conditions (28c) and (28d).
3. Obtain the noise decomposition (29a) irrespectively of cell age by performing the average in Eqs. (29b).

Step 1 can be efficiently computed using numerical root-finding methods. The shooting method in Step 2 consists of an iterative procedure by which the mean molecule number, intrinsic variance and extrinsic variance are obtained through an initial guess on their values after cell division, *E_ρ_*[*x|*0], Σ_int_(0) and Σ_∝_(0), and the result is then refined using standard root-finding methods until the boundary conditions (28c) and (28d) are matched. Step 3 is easily carried out alongside the numerical integration of Step 2. The procedure typically evaluates the noise decomposition in seconds on a desktop computer and may therefore be adequate for statistical inference.

##### a. Suppressing intrinsic or extrinsic noise through feedback mechanisms

Over 40% of known transcription factors in *E. coli* regulate their own expression^40^. We here investigate the sensitivity of negative autoregulatory feedback to cell cycle fluctuations. We consider transcription and degradation of mRNA molecules from which proteins are synthesised

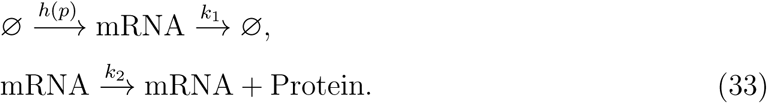

The effect of negative feedback is modelled via a Hill-function for the transcription rate 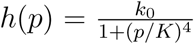, which decreases with the protein number *p*. This approximation is appropriate when the promoter-binding is extremely fast^41^.

In Fig. 4a we show mRNA levels in lineages decrease with cell cycle variability for various feedback strength (the inverse of the dissociation constant, 1/*K*). Mean mRNA numbers in the snapshot statistic either decrease (weak, moderate feedback) or increase with cell cycle variability (strong feedback) depending on the feedback strength. In contrast, protein levels increase with cell cycle variability in lineages but decrease in the snapshot for various feedback strengths (Fig. 4b). In agreement with this trend, we find that intrinsic noise always increases with cell cycle variability while the opposite holds for weak to moderate feedback but not for strong feedback (Fig. 4d). Strikingly, due to the negative feedback regulation, the sensitivity of intrinsic noise of mRNAs is the opposite (Fig. 4c). In contrast to the intrinsic noise properties of the circuit, the total extrinsic noise of the circuit always increases with cell cycle variability, both in lineages and snapshots (Fig. 4d,e). In all cases, our approximations are in good agreement with exact stochastic simulations (Fig. 4 dots) carried out using the First-Division Algorithm given in Ref.^31^ including two non-interacting reporter networks.

**FIG. 4.**
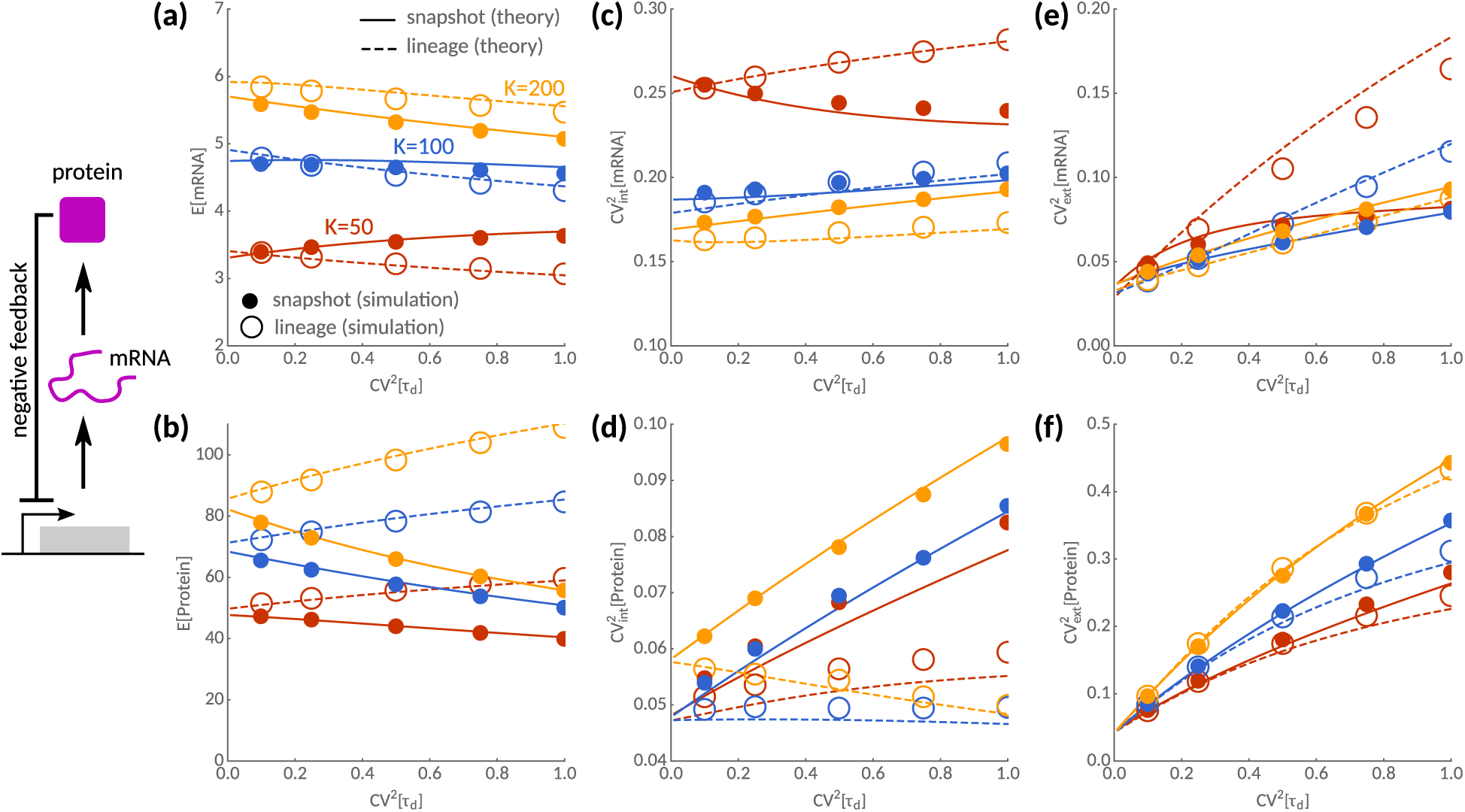
Noise decomposition of a negative feedback circuit. Sensitivities to cell cycle noise CV^2^[*τ_d_*] of mean, intrinsic and extrinsic noise contributions are shown for weak (yellow, *K* = 200), moderate (blue, *K* = 100) and strong feedback (red, *K* = 50). Predictions by the linear noise approximation (solid lines) are in good qualitative agreement with stochastic simulations (dots). (**a**) In lineages, the mean mRNA number always decreases with cell cycle variability while this is not true in snapshots for moderate to high feedback. (**b**) In contrast, protein levels always increase in lineages but decrease in snapshots. (**c**) The corresponding intrinsic noise profiles of mRNAs typically increase with cell cycle noise except in snapshots with strong feedback. (**d**) Intrinsic noise of proteins always increases with cell cycle noise in snapshots but not in lineages. (**e,f**) Total extrinsic noise increases with cell cycle variability for mRNAs and proteins. However, strong feedback may significantly reduce extrinsic noise in snapshots compared to lineages. Deviations between the approximation (lines) and the simulations (dots) are most pronounced for strong feedback. Parameters are *k*_0_ = 10, *k_m_* = 1, *k_s_* = 10 and unit-mean interdivision times.

Finally, we use the noise decomposition to understand how heterogeneity can be controlled by natural and synthetic circuits. Negative feedback is widely known to reduce noise but often requires fine-tuned parameters^42–44^. How this translates to individual functional noise components, such as intrinsic and extrinsic noise, has only been explored in response to parameter fluctuations^6,45^ but not in the context of the ubiquitous population dynamics. Here, we are specifically interested in the sensitivity of lineage and population snapshot statistics to cell cycle noise.

In Fig. 5a we show that negative feedback can efficiently suppress intrinsic noise as the feedback strength is varied. Intriguingly, comparing the minimum noise levels in lineages and snapshots, vastly different values of the dissociation constants achieve noise suppression in these measures. To study this dependency in more detail, we compute the optimal feedback strength that minimises intrinsic noise as shown in Fig. 5b. Intriguingly, the optimal values exhibit opposite sensitivities to the cell cycle variability in lineages than in the population snapshots. To efficiently suppress intrinsic noise in a lineage, we must decrease the feedback strength in response to an increase in cell cycle variability. To compensate for intrinsic variability across the population, however, the feedback strength must increase by almost a two-fold of what would be required in the lineage.

**FIG. 5.**
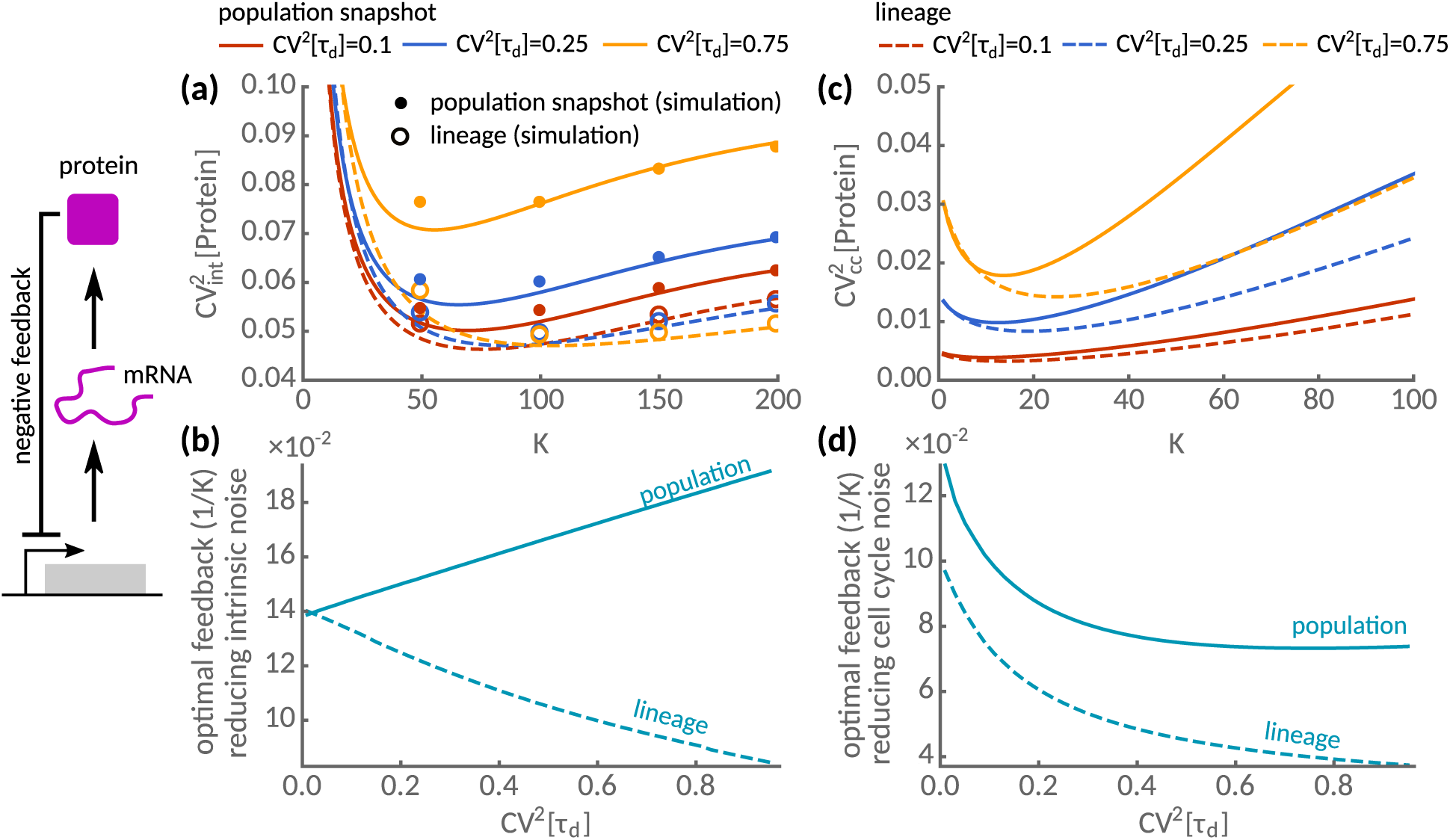
Feedback strategies for noise suppression in lineages and populations. Intrinsic and extrinsic noise statistics of negative autoregulatory feedback circuit are shown as a function of *K*, the inverse feedback strength, for three different levels of cell cycle noise CV^2^[*τ_d_*] = 0.1 (red), 0.25 (blue) and 0.75 (yellow). (a) Intrinsic noise exhibits a minimum as a function of the repression strength both in lineage (dashed) and in snapshot statistics (solid lines). The predictions obtained using the linear noise approximation (lines) are in good agreement with exact stochastic simulations using the First-Division Algorithm^31^ (dots for population, open circles for lineages). (b) Optimal feedback strength (1/*K*) to minimise intrinsic noise is shown. The feedback strength increases with interdivision time noise in lineages but decreases in population snapshots. (c) The transmitted cell cycle noise shows a minimum in dependence of the repression strength both in lineage (dashed) and in snapshot statistics (solid lines). (d) The optimal feedback strength to minimise transmitted cell cycle noise decreases with division time noise both in lineages and to a lesser extent in the population. Parameters as in Fig. 4.

In other situations, it may be advantageous to reduce the extrinsic instead of the intrinsic noise component. In Fig. 5c, we show that tuning the dissociation constant (*K*) can similarly reduce the transmitted cell cycle noise 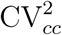. Comparing the optimal feedback strength (1/*K*) as a function of the cell cycle noise CV[*τ_d_*] (Fig. 5d), we observe that they increase with cell cycle variability in both lineage and snapshot. It is worth noting that vastly different feedback strengths achieve either intrinsic or extrinsic noise suppression (cf. Fig. 5c,d). These findings highlight that a single feedback loop may not be su cient to simultaneously suppress both noise components whether in lineages or population snapshots.

## IV Discussion

We presented an analytical framework to analyse stochastic biochemical reactions in an exponentially growing cell population. This theory allows us to characterise and systematically decomposes cellular noise into intrinsic and extrinsic components, which applies to general stochastic biochemical networks. We found that a typical cell in the population expresses lower levels of proteins per cell than an isolated cell tracked over successive cell divisions. As a consequence, we observed higher levels of intrinsic noise but, for the examples studied, the extrinsic noise component was significantly reduced. These effects are most pronounced in the presence of division time variability as it is the case in natural populations. Importantly, this highlights that one needs to account for cell cycle fluctuations when modelling either intrinsic or extrinsic noise components.

Previous studies^46,47^ focussed solely on the effect of age-structure but mostly neglected cell cycle variations. We demonstrated that the statistics of lineages and population snapshots are not equivalent even when the cell cycle position is known. Although these differences appear to be small when divisions occur deterministically, they will be pronounced in the presence of division time variability (Figs. 3 and 4). In particular, we showed that measuring cells within a narrow range of cell cycle stages, as for instance achieved through gating^8,28^, does not eliminate all extrinsic noise due to cell cycle fluctuations. In reality, cells are affected by more than one type of extrinsic noise source as reaction rates may fluctuate over time and between cells^6,7,13^. These effects should be added to the transmitted extrinsic noise. We anticipate, however, that it will be difficult in practice to distinguish these fluctuations from the variations induced by cell cycle variability.

While this study focused on the effects of age-structure on biochemical dynamics, several simulation studies suggest that cell size also coordinates gene expression^48,49^. Incorporating additional physiological details such as cell size into our framework could thus provide insights to the statistics of intracellular concentrations^50–52^ and extrinsic noise transduced from cell size control and growth rate fluctuations^53^. A different limitation of this study is that it is based on the linear noise approximation, which albeit being exact for linear reaction networks, represents an approximation assuming large molecule numbers. Its estimates can be inaccurate for nonlinear reaction networks involving low numbers of molecules. An improvement to this approximation could employ higher order terms in the system size expansion^41,54^, or close the hierarchy of moments using moment closure approximations^55^.

Heterogeneity inferred from snapshots is often used to say something about a cell’s history. By grouping cells of similar ages, as in ergodic rate analysis^56^, one can in principle reconstruct time-course information. We demonstrated that such a procedure produces different results to the lineage statistic (see Fig. 2). Instead, the variability across the population is equivalent to choosing an arbitrary cell from the final population and tracing it backwards in time^31^. Although this equivalence provides a sample-path interpretation of snapshot data, it is worth pointing out that it does not apply when cell ages are unidentified. In this case, understanding the relationship between single cell fluctuations and population heterogeneity requires an agent-based framework as the one presented.

We showed that gene expression noise in populations is coupled to the population growth rate, as observed in population studies^57^. This dependency is crucial when quantifying summary statistics such as mean and variances. We found that the cellular heterogeneity displays opposite sensitivities to cell cycle variability across populations and lineages. For negative feedback circuits, this implies that no parameter tuning enables cells to minimise noise of both measures efficiently (Fig. 5). Reducing noise in lineages over time comes at the cost of increased population heterogeneity, a strategy cells could exploit to diversify in response to stress^58^. Conversely, tuning snapshot homogeneity sacrifices lineage-optimality, which could confer advantages when gene expression couples to global physiological factors such as cell size, growth rate or cell division^59^.

Cells may thus perform suboptimally depending on which experimental setup is used to study them, whether it is a mother machine or a chemostat. Identifying the relevant noise components and cellular objectives will likely depend on the environmental and experimental conditions, or even on the particular application^58,59^. These dependencies thus reveal a fundamental trade-off for the evolution of natural circuits and the design of synthetic circuits in living cells.

In summary, we presented an agent-based framework for the statistical analysis of population snapshots. Inherent in this approach are several noise sources that reveal typical features of snapshot data using noise decompositions. The present framework is widely applicable and as such it also applies to large gene regulatory, signalling or metabolic networks. We, therefore, envision that the proposed moment-based approach could prove especially useful for parameter inference from snapshots of living cells^60^.

## V Supporting information

### A Statistics of interdivision times and age distributions

We here characterise the statistics of division times and age-distributions in lineages and populations. To this end, it is useful to recall the definition of the Laplace transform of the interdivision time distribution *φ*(*τ_d_*) in a lineage

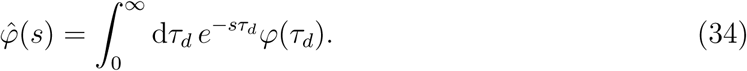

We assume that either *φ*(*τ_d_*) is known or can be calculated from the division rate via Eq. (7).

#### 1 Moments of interdivision times in a population

To calculate the statistics of the interdivision times in the population, we employ the Laplace transform of Eq. (11), which is

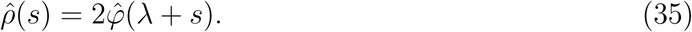

and note that 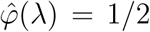 due to characteristic equation (8). The moments can thus be expressed in terms of the Laplace transform

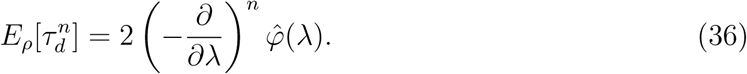

From these, we can compute

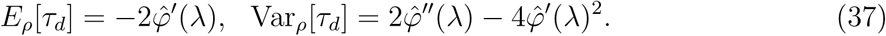

and hence

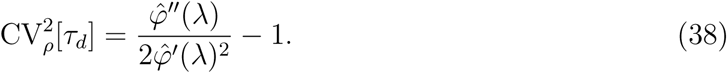

#### 2 Age-distribution in lineages

The age-distribution yields the frequency of cell ages observed for different single cell measures. To compute the moments of the age-distribution in a lineage, we compute the Laplace transform of Eq. (13), which gives

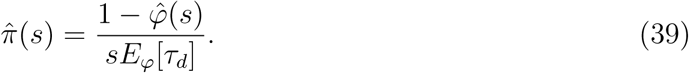

By differentiating the above expression repeatedly at *s* = 0, we find

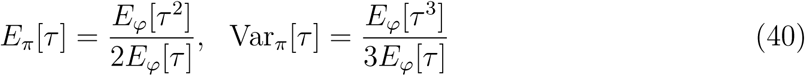

and

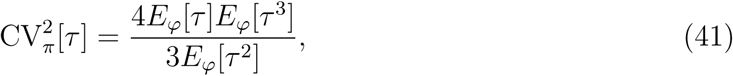

which concludes the the age-statistics in the population snapshot.

#### 3 Age-distribution in populations

Similarly, we consider the Laplace transform of the age-distribution in a population snapshot, Eq. (6), which evaluates to

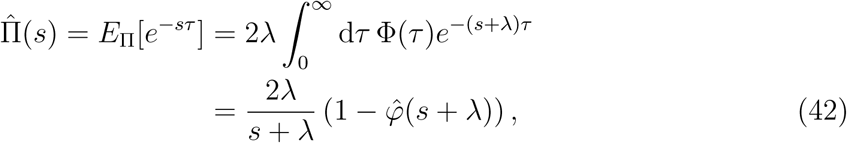

Repeated differentiation at *s* = 0, gives

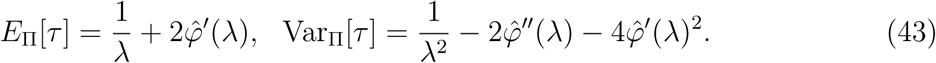

It is straightforward to evaluate these statistics numerically as we do for the log-normal distribution in Fig. 2 and 3. For the gamma-distribution, the population growth rate, the age- and interdivision-time distributions can be obtained in close form as we show in the following.

#### 4 Gamma distribution: Explicit solutions to population growth rate and the age/interdivision time distributions

We fix the division time to be gamma distributed with density function

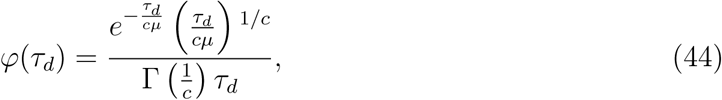

where Γ is the gamma function, such that *E_φ_*[*τ_d_*] = *μ* and 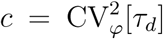. The Laplace transform of the distribution is

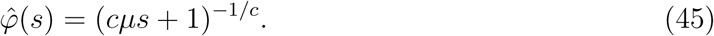

Recasting now the Euler-Lotka equation in the form 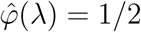, we can solve for *λ* to obtain

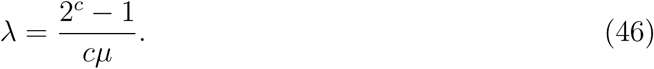

The division time distribution in the population is then given by

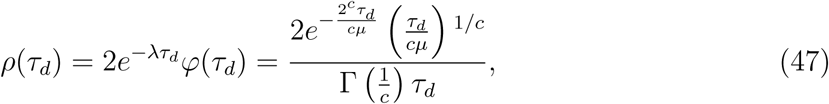

which is also a gamma distribution but with a shorter mean division time but the same coefficient of variation

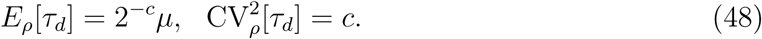

Further, the age-distribution in a lineage is

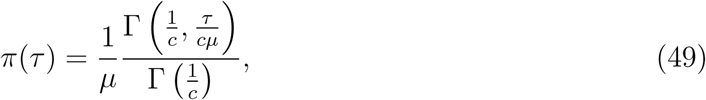

where Γ(*·, ·*) is the upper incomplete gamma function. Its statistics are with statistics

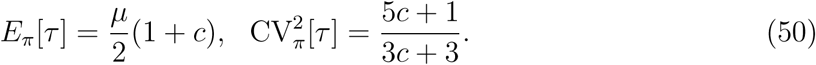

Similarly, the age-distribution in the population becomes

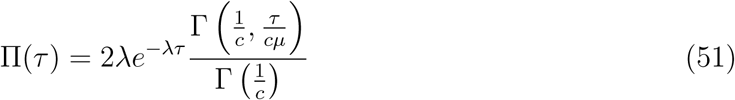

with statistics

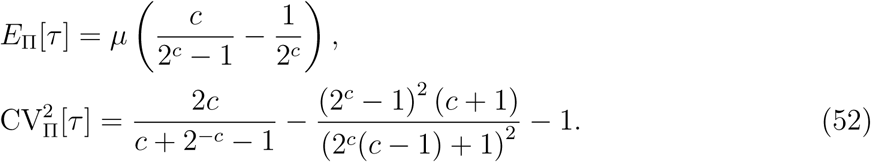

Interestingly, it follows that *E*_Π_[*τ*] < *E_Π_*[*τ*], but 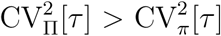 for *c* < 1 and 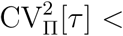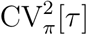 for *c* > 1.

### B An explict formula for the uncertainty due to unknown cell age

We here verify Eq. (31c) of the main text, which holds for linear reaction networks. To this end we define ϵ (*τ*) = *E*_Π_[*x|τ*] − *E*_Π_[*x*] such that 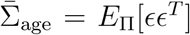 and use Eq. (18a) to write

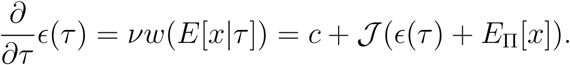

We used the fact that *vw*(*E*[*x|τ*]) = *c* + 𝒯 *E*[*x|τ*], where *c* is a constant vector, since for linear reaction networks the propensities are linear in the number of molecules. Making use of Eq. (53) we then compute

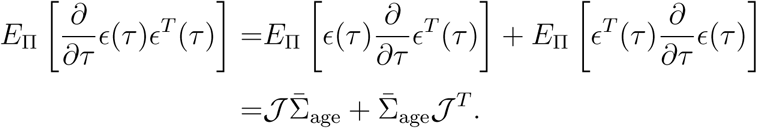

On the other hand, using Eq. (15) of the main text, it follows that

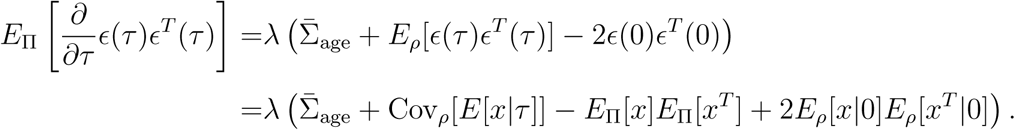

Combining the last two equation gives the result (31c) of the main text.

### C Detailed discussion of the moment-closure conditions

While the moment equations derived in Sec. II B are exact, the equations for cells of the same age are only closed when *w_r_*(*x*) depends at most linearly on the molecule numbers *x* and the covariance of the partitioning kernel Cov*_B_*[*x|x′*] depends at most quadratically on the number molecules in the mother cell *x′*. This holds, for instance, for biochemical composed solely from unimolecular reactions and independent binomial partitioning. Similarly, it holds true for the mean of cells with unknown age, but not generally for their corresponding variances. Specifically, the covariance for cells of unknown age also depends on the moments for cells of known age and thus they must explicitly depend on the division time distribution.

There are now two scenarios in which the variances are independent of the division time distribution. The first case is when the age-distribution coincides with the division time distribution Π(*τ*) = *ρ*(*τ*), which follows only when the division rate *γ* is constant and independent of age, i.e. the division times are exponentially distributed. The second case assumes a particular division kernel *B*(*x|x′*) that satisfies 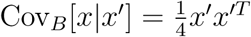, which follows when all molecules are inherited by only one of the daughter cells. In all other cases, which seem most relevant in practice, the moment equations for unknown cell age involve the moments for cells of known age. Thus, for general nonlinear reaction networks, they involve two hierarchies of moments that cannot be easily closed. A simple and generally applicable approximation that circumvents this problem is given in Sec. III A using the linear noise approximation.

### D Analytical noise decomposition for gene expression with degradation

We consider a simple system in which a protein *P* is translated in stochastic bursts of size *m* and is subsequently degraded

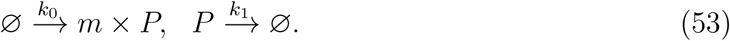

Because the burst size is a random variable we can recast the synthesis reaction into a series of reactions with reaction rates *k*_0_*π*(*m*),

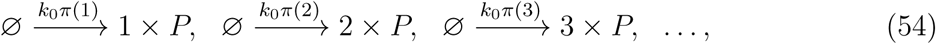

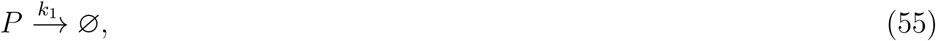

where *π*(*m*) is the distribution of burst sizes, which is geometric for the two-stage model of gene expression.

#### 1 Mean protein number of the same age

The equation for the mean number of molecules is then

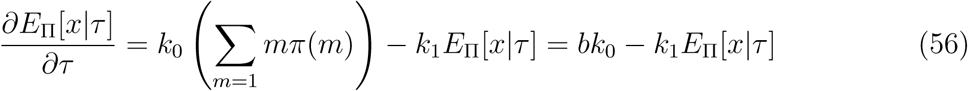

with solution

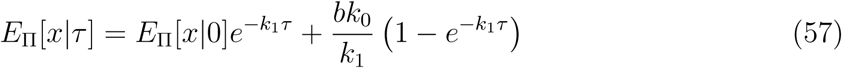

Substituting the solution into the boundary condition *E_ρ_*(*E*_Π_[*x|τ*]) = 2*E*_Π_[*x|*0] and solving for *E*_Π_[*x|*0] yields the final result

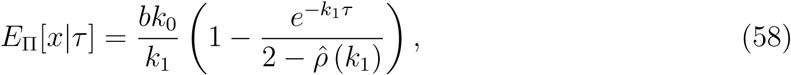

where 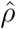 is the Laplace transform of the division time distribution

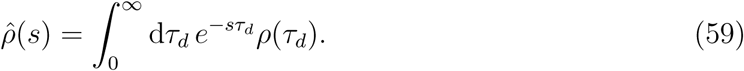

As we will show in the following the statistics of gene expression in growing populations depends crucially on this function.

#### 2 Protein fluctuations for cells of the same age

To compute the protein fluctuations, we see from (58), that the Jacobian is

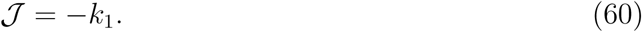

The diffusion matrix then follows from Eq. (19) then follows

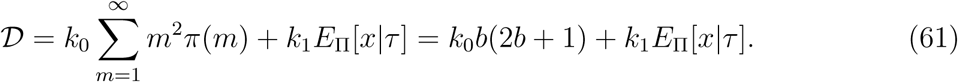

The variance of intrinsic and extrinsic fluctuations obeys Eqs. (18b), which read explicitly

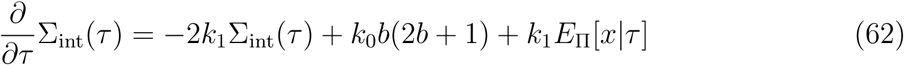

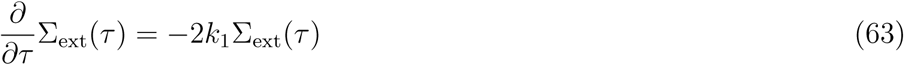

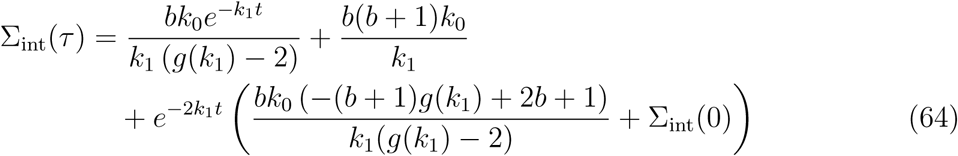

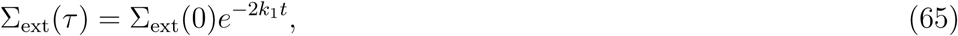

where Σ_int_(0) and Σ_ext_(0) are the intrinsic and extrinsic variation at cell division, which have to be determined from the boundary conditions. According to Eq. (28c) and (28d), the boundary conditions are

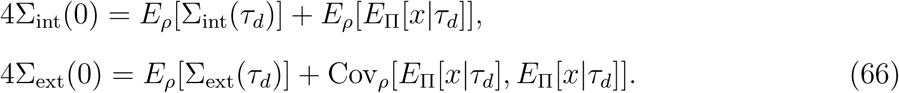

To compute these values we notice that the variances at cell division follow from averaging Eqs. (64) over the division time distribution *ρ*, which results in

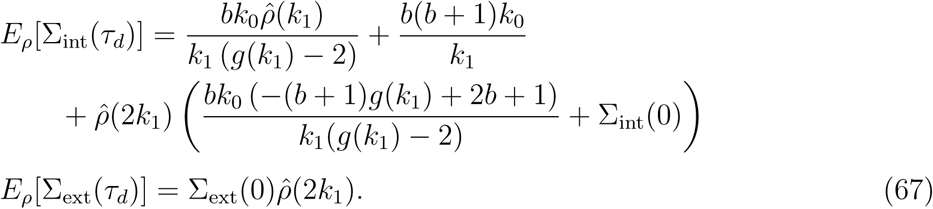

Further, we evaluate

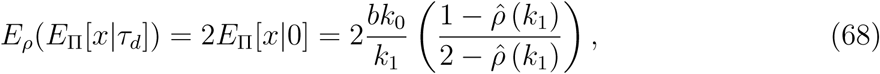

and

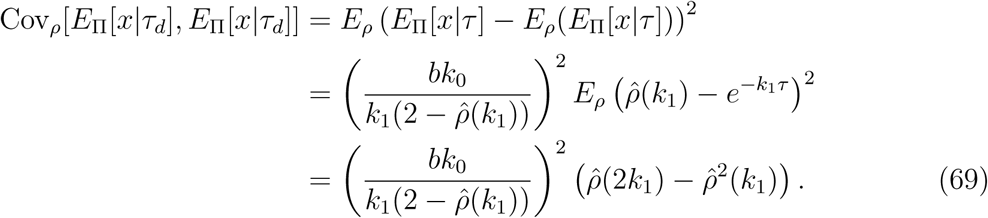

Plugging Eqs. (67), (68) and (69) into (66), solving for Σ_int_(0) and Σ_ext_(0) and using the result in Eqs. (64), we finally arrive at

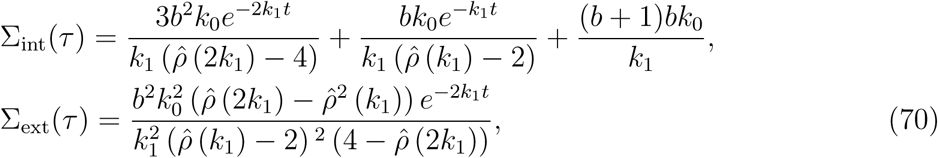

which determines the progression of intrinsic and extrinsic fluctuations over the cell cycle.

#### 3 Protein statistics for cells of unknown age

The mean protein number is given by

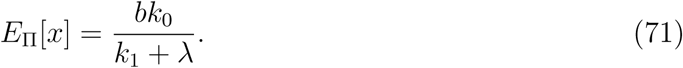

Thus the mean number is determined from the balance between the rates of translation, degradation and dilution due to cell divisions. From Eqs. (31) we compute

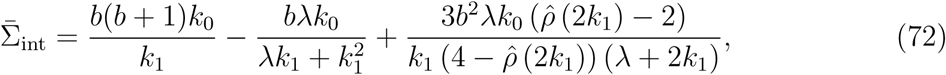

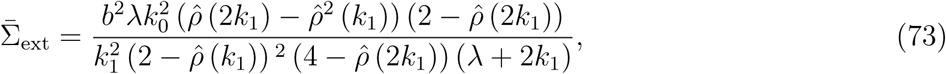

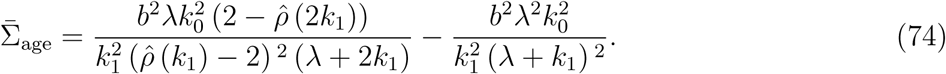

Finally, we compute 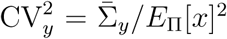 to arrive at the expressions for the coefficient of variations

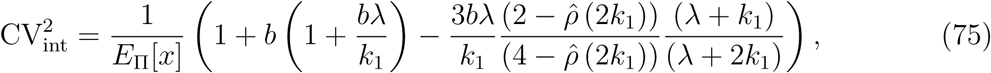

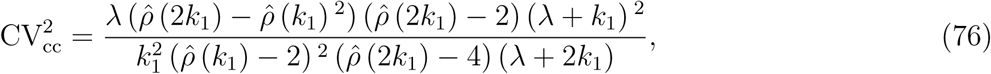

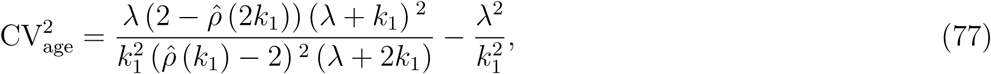

which denote the intrinsic noise, the transmitted noise from cell cycle fluctuations and the uncertainty due to the unknown cell age. It is obvious that these expressions are much more involved than for the case without degradation because they depend on the Laplace transform 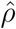 of the division time distribution. Interestingly, the last two components, which together represent the extrinsic noise, also depend on the degradation rate *k*_1_ meaning that the total extrinsic noise is constant only when measured the mean concentration is varied through the either transcription rate or burst size.

## Acknowledgements

Acknowledgments

It is a pleasure to thank Vahid Shahrezaei for valuable feedback.

